## Supplementary Figures 1-10 for "Continuous, Topologically Guided Protein Crystallization Controls Bacterial Surface Layer Self-Assembly"

Supplementary Information (Extended Data)

### Continuous, Topologically Guided Protein Crystallization Controls Bacterial Surface Layer Self-Assembly

Colin J. Comerci*[1,2], Jonathan Herrmann*[3,4], Joshua Yoon[5,2], Fatemeh Jabbarpour[3,4], Xiaofeng Zhou[6], John F. Nomellini[7], John Smit[7], Lucy Shapiro[6], Soichi Wakatsuki[3,4], and W.E. Moerner[1,2,5]

[1]Biophysics Program, Stanford University, Stanford, CA USA

[2]Department of Chemistry, Stanford University, Stanford, CA USA

[3]Department of Structural Biology, Stanford University, Stanford, CA USA

[4]Bioscience Division, SLAC National Accelerator Laboratory, Menlo Park, CA USA

[5]Department of Applied Physics, Stanford University, Stanford, CA USA

[6]Department of Developmental Biology, Stanford University, Stanford, CA, USA

[7]Department of Microbiology and Immunology, University of British Columbia, Vancouver, BC, CA

*These authors contributed equally to this work.

**Supplementary Figure 1**

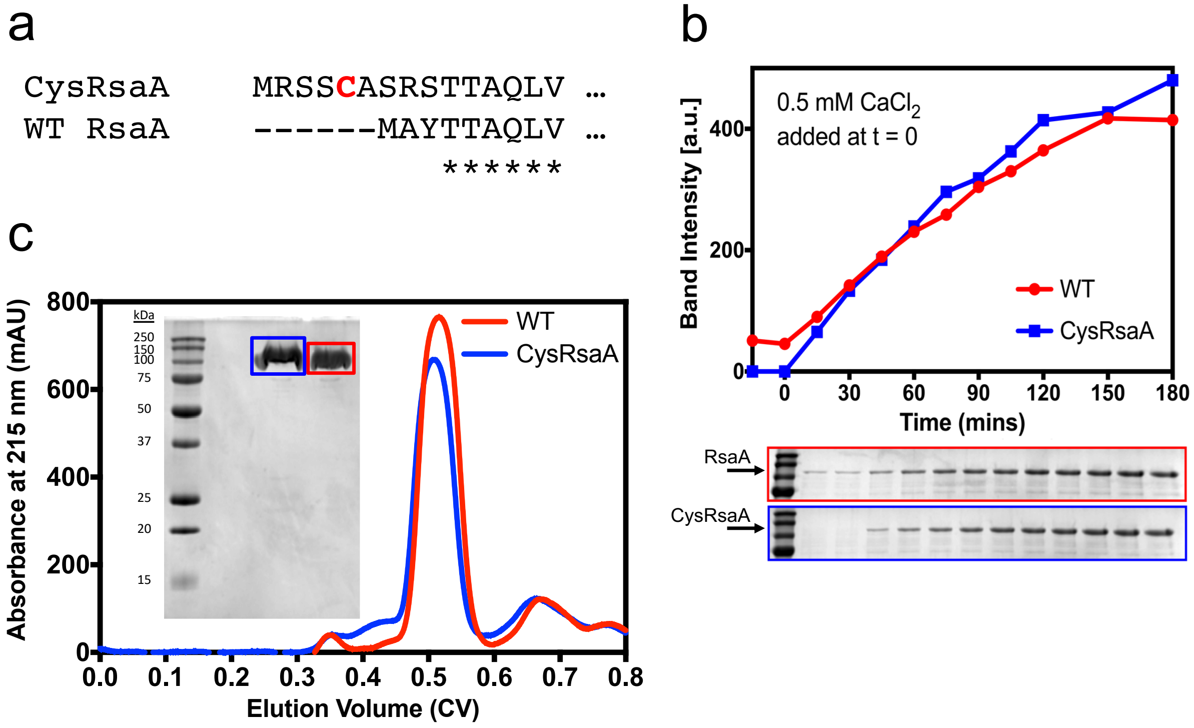


**Supp Fig 1.** CysRsaA cells produce native-like S-layer protein. a) Amino acid sequences of the N-termini of WT and modified (CysRsaA) RsaA protein. The cysteine used for fluorescent labeling is shown in bold red. b) Time-resolved blot showing the appearance of soluble S-layer protein upon calcium addition. CysRsaA protein is produced at a similar rate to WT RsaA. c) Size exclusion chromatograms of CysRsaA and WT RsaA protein on an S200 size exclusion column. Inset: Coomassie-stained SDS-PAGE of purified CysRsaA and WT RsaA protein.

**Supplementary Figure 2**

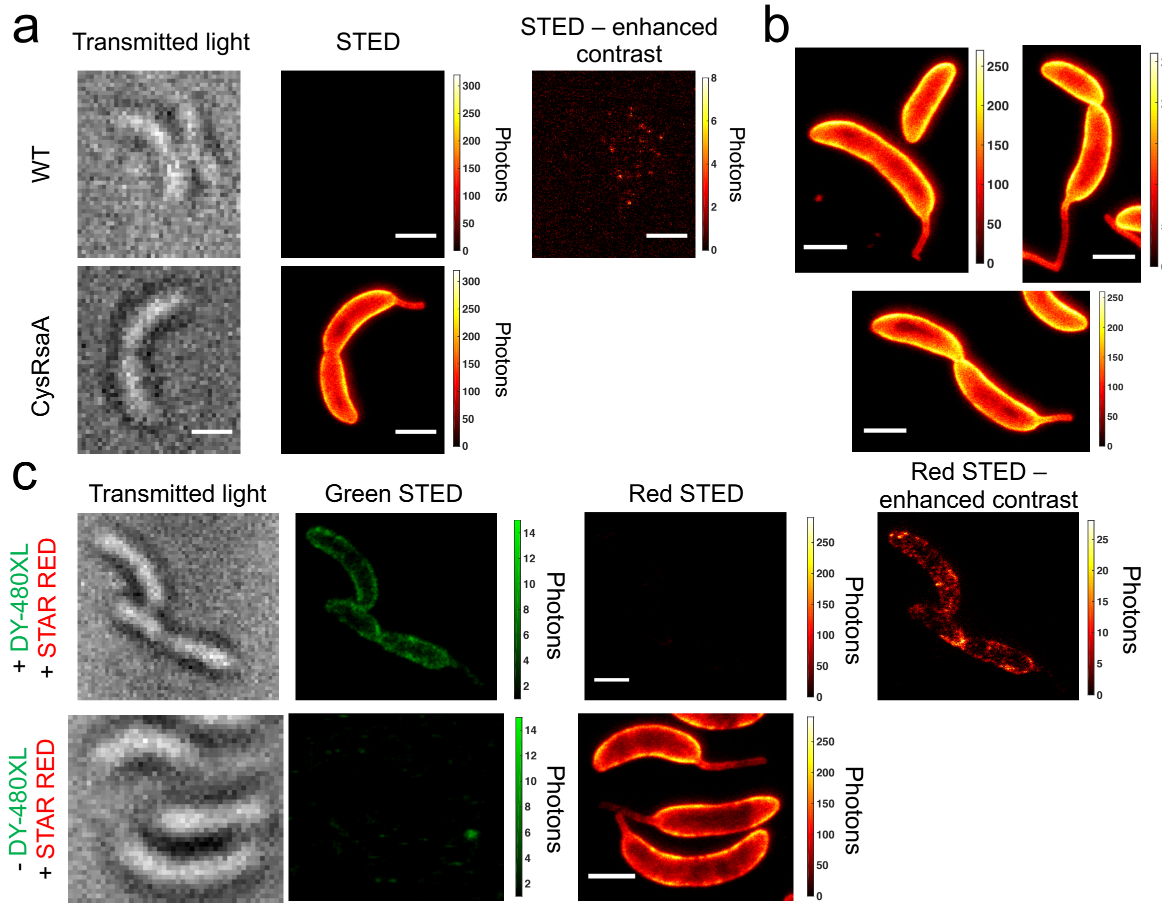


**Supp Fig 2.** Maleimide chemistry provides a highly efficient method to specifically label the CysRsaA S-layer on living *Caulobacter crescentus* cells. a) WT cells (top) show almost no labeling upon incubation with STAR RED-maleimide, while CysRsaA cells (bottom) show clear and complete labeling of the cell surface. b) Additional representative CysRsaA cells labeled with STAR RED showing complete labeling. c) Top: CysRsaA cells labeled first with DY-480XL-maleimide (green) followed immediately by STAR RED-maleimide (red) show very little labeling by STAR RED. Bottom: cells labeled only with STAR Red-maleimide (bottom) show uniform labeling, demonstrating that labeling with DY-480XL efficiently labels and blocks CysRsaA sites present on the cell surface. Scale bars = 1 µm.

**Supplementary Figure 3**

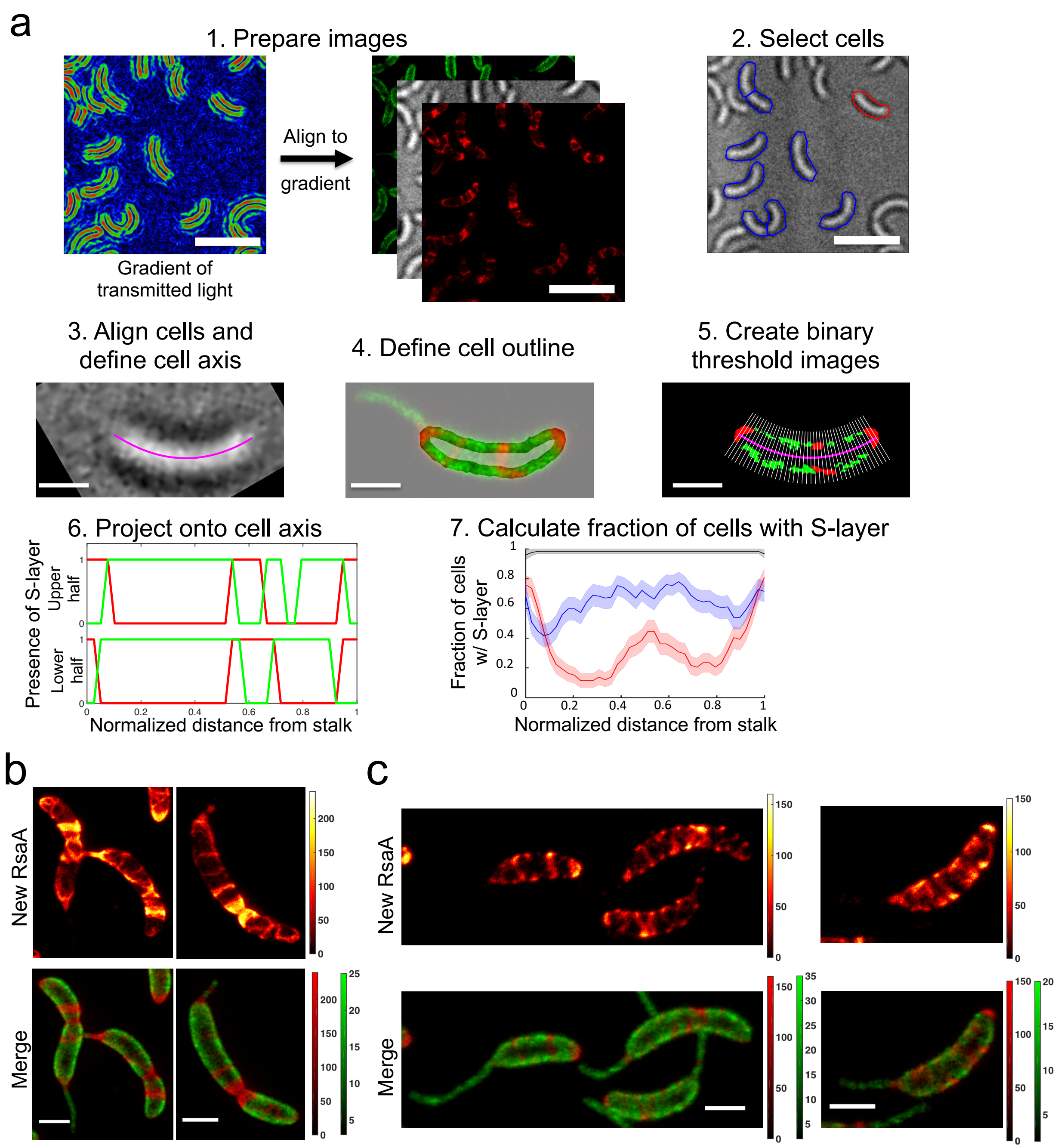


**Supp Fig 3.** In-depth schematic of binary cell profile analysis algorithm as described in Methods. a) 1. Images are prepared for analysis by aligning the fluorescent images to the gradient of the transmitted light image (left). 2. Cells are selected by the user drawing an outline around the cell periphery. Red outlined cell is shown in 3-6. 3. The cells are aligned to be horizontal, using a radon transform, with the stalk on the left-hand side. The cell axis (magenta line) is defined by fitting a $2^{nd}$ order polynomial to the maximum transmitted light pixels. 4. A mask of the cell outline is used to select the analyzed area (transparent in image). 5. A binary image is created in each color channel (red and green) by thresholding the fluorescence images.

The positive pixels are projected onto the cell axis (magenta line) and binned using 40 equi-length bins (thin white lines). 6. The projected binary images create 2 binary cell profiles, for the upper and lower half of the cell (top and bottom), in each color channel (red and green lines). 7. The upper and lower binary cell profiles from many cells are used to calculate the probability of finding new S-layer at a given normalized distance along the cell. Plot shows new S-layer for control (black), native S-layer production (red), and exogenously added purified CysRsaA (blue), recreated from Fig 1j. b) Additional representative cells showing native S-layer production. c) Additional representative cells showing exogenous addition of purified CysRsaA to ΔRsaA cells. Scale bars = 5 µm (panels a1-a2); 1 µm (panels a3-a5; b and c).

**Supplementary Figure 4**

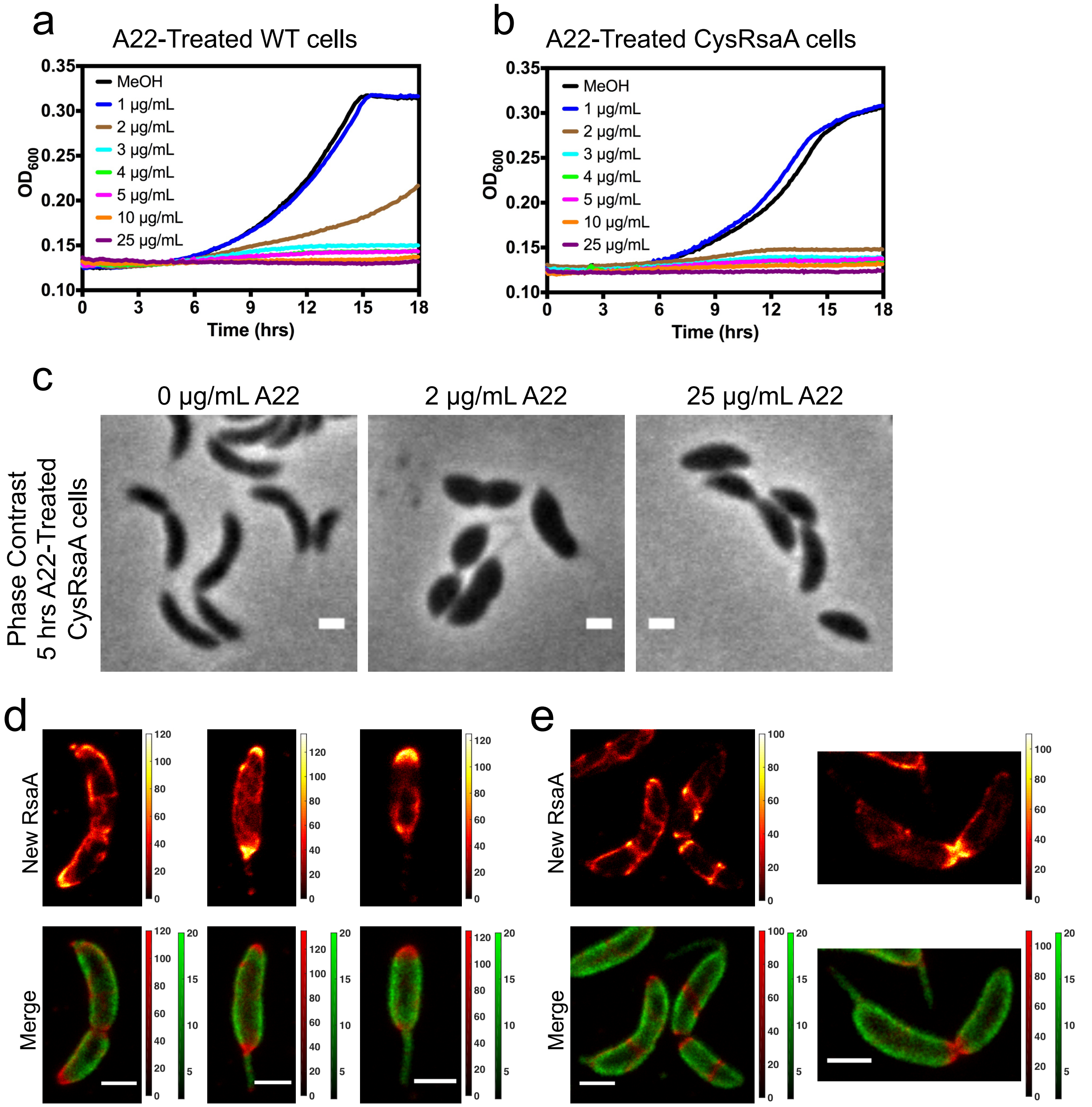


**Supp Fig 4.** A22 affects WT and CysRsaA cells similarly, and cell growth is concentration dependent. Growth curves are shown of a) WT cells or b) CysRsaA cells grown in varying concentrations of A22. Significant growth inhibition begins to occur at 2 μg/mL A22 while 25 ug/mL A22 stops growth entirely. c) CysRsaA cells treated with the stated concentration of A22 for 5 hrs. 2 μg/mL A22 leads to lemon-shaped cells, illustrative of delocalized PG insertion. Cell viability with 2 μg/mL A22 as measured by colony forming units arising from equal volumes of treated and untreated cells was 100 ± 2% for WT cells and 99 ± 2% for CysRsaA cells (n=3). 25 μg/mL A22 stops cell growth, leading to cells more similarly shaped to untreated cells. Cell viability with 25 μg/mL A22 was 56 ± 9% for WT cells and 57 ± 7% for CysRsaA cells (n=3). Additional representative cells showing native S-layer production on cells treated with 2 μg/mL A22 (d) or 25 μg/mL A22 (e). Scale bars = 1 μm.

**Supplemental Figure 5**

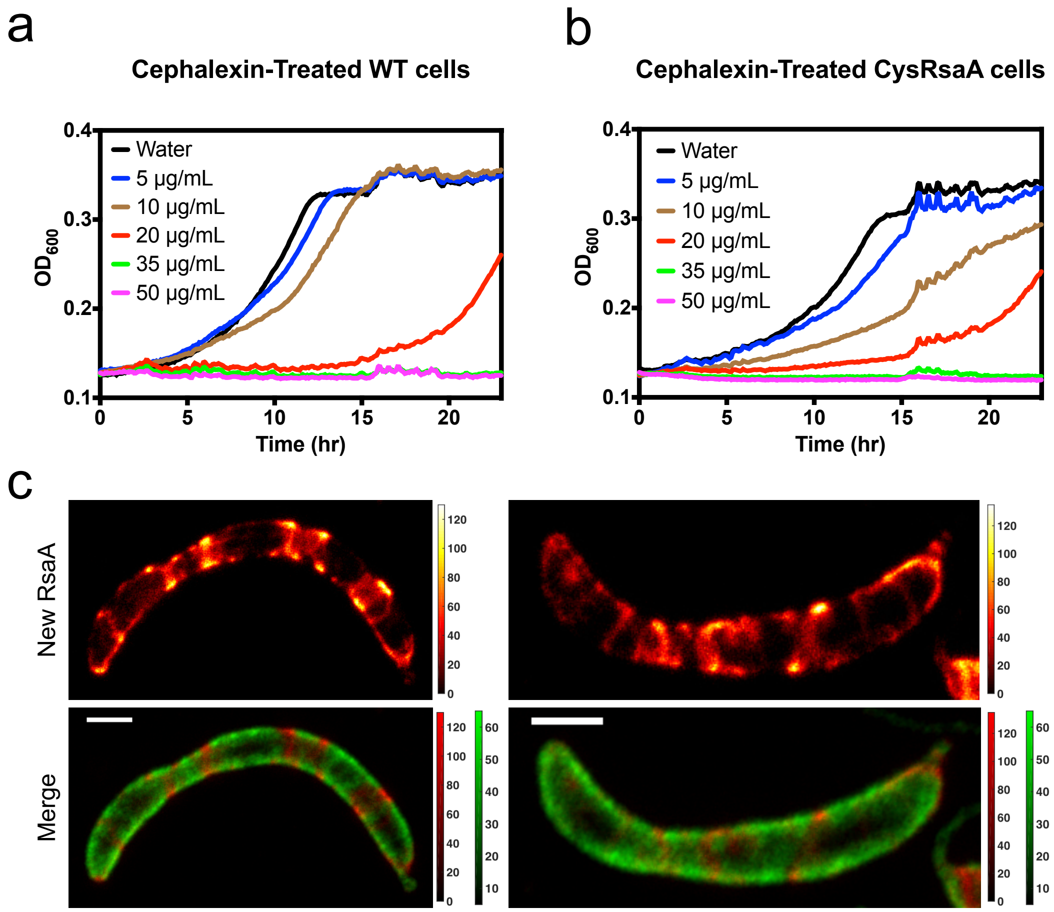


**Supp Fig 5.** Cephalexin affects WT and CysRsaA cells similarly. Growth curves are shown of a) WT cells or b) CysRsaA cells grown in varying concentrations of cephalexin. Significant growth inhibition begins to occur at 5-10 µg/mL cephalexin while 35 ug/mL cephalexin stops growth entirely. c) Additional representative cells showing native S-layer production on cells treated with 35 µg/mL cephalexin. Scale bars = 1 µm.

Supplementary information to Comerci and Herrmann *et al*, "Continuous, Topologically Guided Protein Crystallization Controls Bacterial Surface Layer Self-Assembly"

**Supplemental Figure 6**

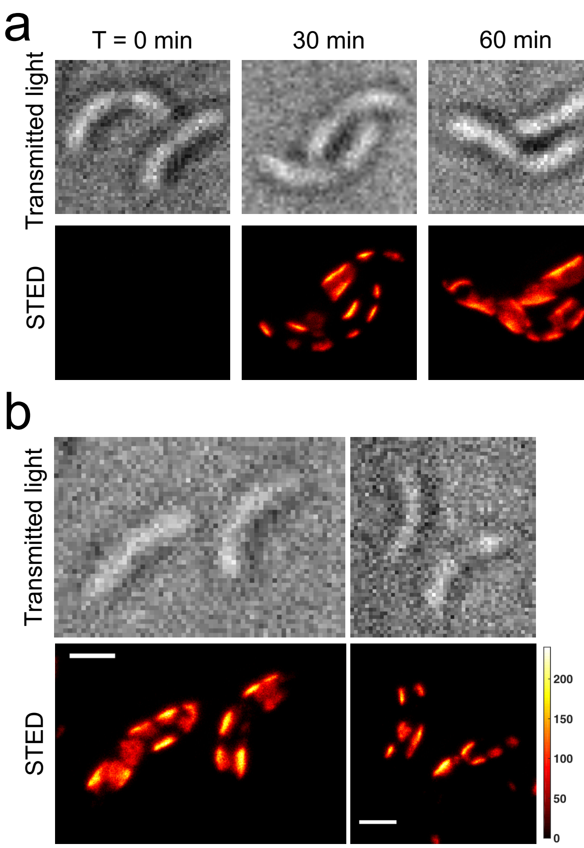


**Supp Fig 6.** Complete time course of *de novo* S-layer growth. CysRsaA cells grown in M2G with 50 µM $CaCl_2$ are resuspended in M2G with 500 µM $CaCl_2$, initiating the *de novo* regrowth of their S-layer. STED imaging reveals the S-layer assembles at discrete patches that grow larger over time, nearly covering the cell by ~2 hours. b) Additional representative cells showing *de novo* S-layer production after 30 minutes. Scale bars = 1 µm.

**Supplemental Figure 7**

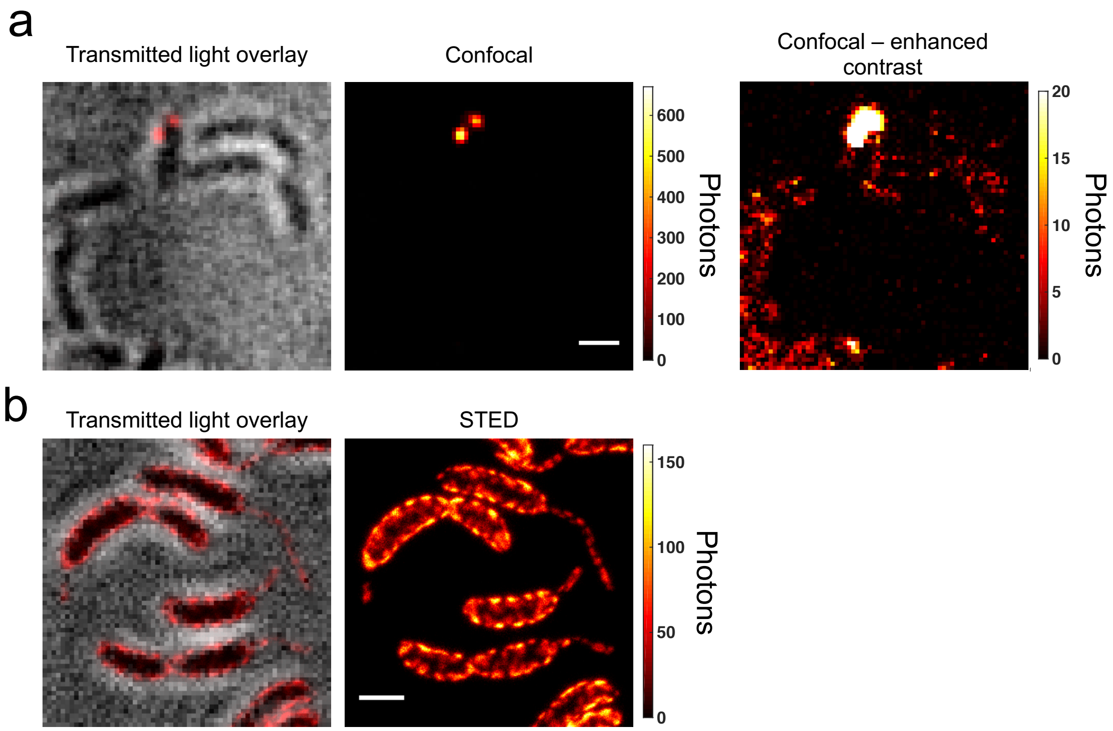


**Supp Fig 7.** Addition of low concentration exogenous CysRsaA suggests diffusing molecules, while addition of high concentrations of exogenous CysRsaA shows creation of many small nucleation puncta. a) Confocal fluorescence image of ΔRsaA cells with 1 nM Cy3 labeled CysRsaA added. Only one cell exhibits crystalline puncta. At high contrast, low fluorescence signal on the cells suggests the presence of diffusing RsaA. b) STED fluorescence images of ΔRsaA cells with 200 nM STAR RED labeled CysRsaA added in one bolus show the presence of many small nucleation puncta. Scale bars = 1 μm.

**Supplemental Figure 8**

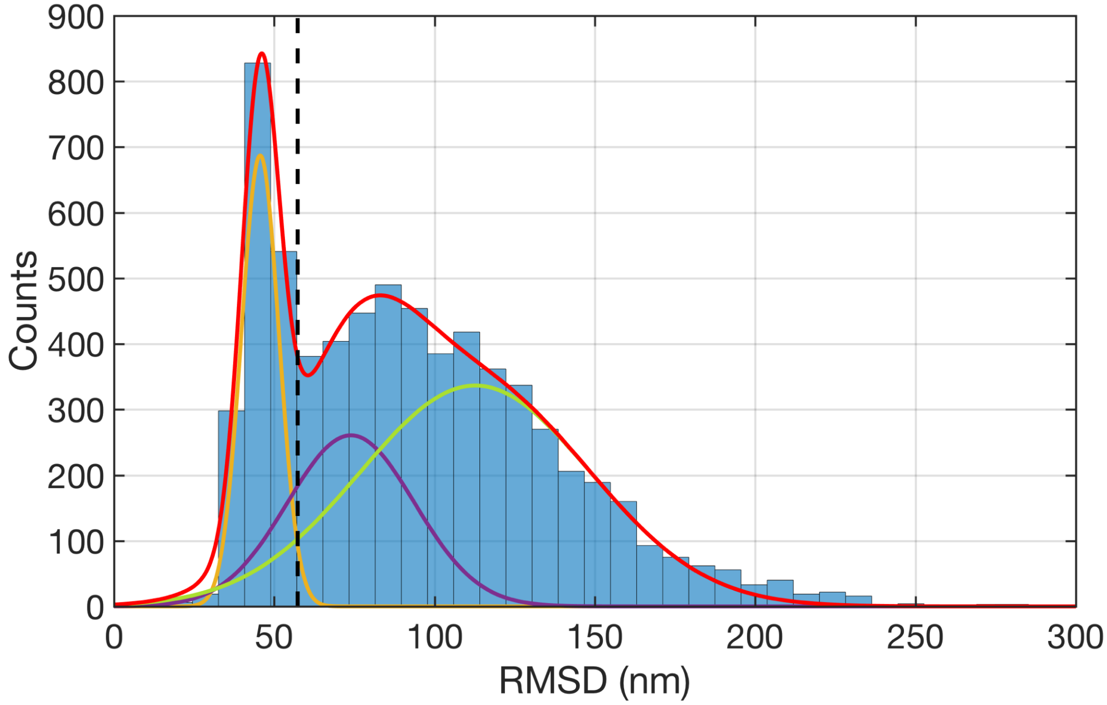


**Supp Fig 8.** Determination of the threshold of the RMSD for bound RsaA molecules. A histogram of all RMSD values from 1 second time windows shows a distinct low RMSD peak corresponding to bound molecules. The distribution was fit to three Gaussians (orange, purple, and green lines), and the threshold of 57.3 nm (vertical black dashed line) was chosen corresponding to 2σ above the mean of the bound Gaussian (orange).

#### Supplemental Figure 9

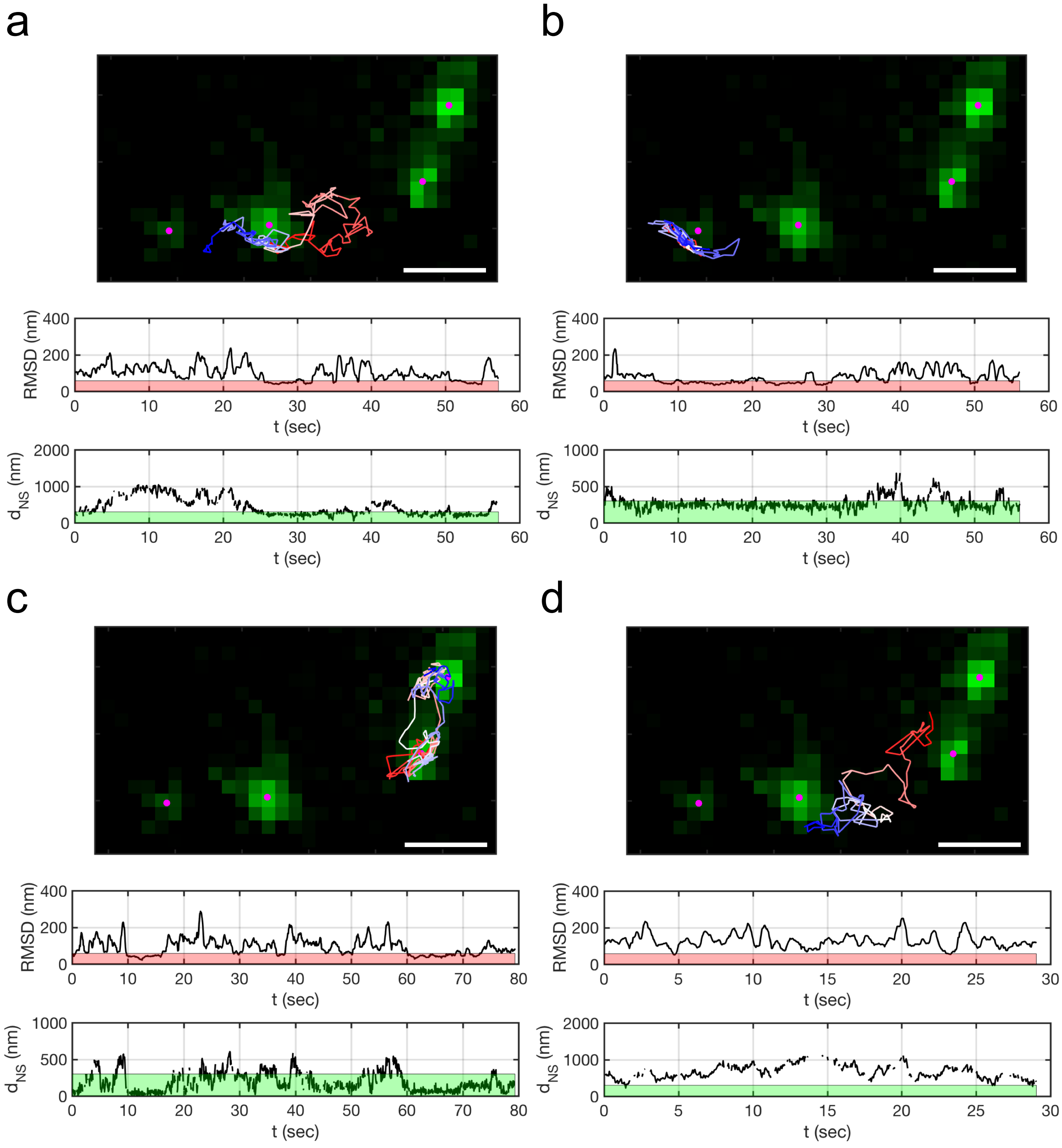


**Supp Fig 9.** 2-color single molecule tracking shows diffusion and binding of RsaA molecules to growing crystals. Four example single particle trajectories (a-d). Upper: tracks are shown in red-to-white-to-blue line, RsaA puncta are shown in green, and the Gaussian-fit centers of the puncta are shown in magenta. Middle: the calculated RMSD for each trajectory as a function of time. The 57.32 nm threshold for determining bound molecules is shown in red. Lower: the distance to the centroid of the closest RsaA punctum. The 300 nm threshold used in Fig 4d is shown in green. Tracks (a-c) show periods of binding as well as diffusion, while (d) shows nearly free diffusion. Scale bars = 1 µm.

**Supplemental Figure 10**

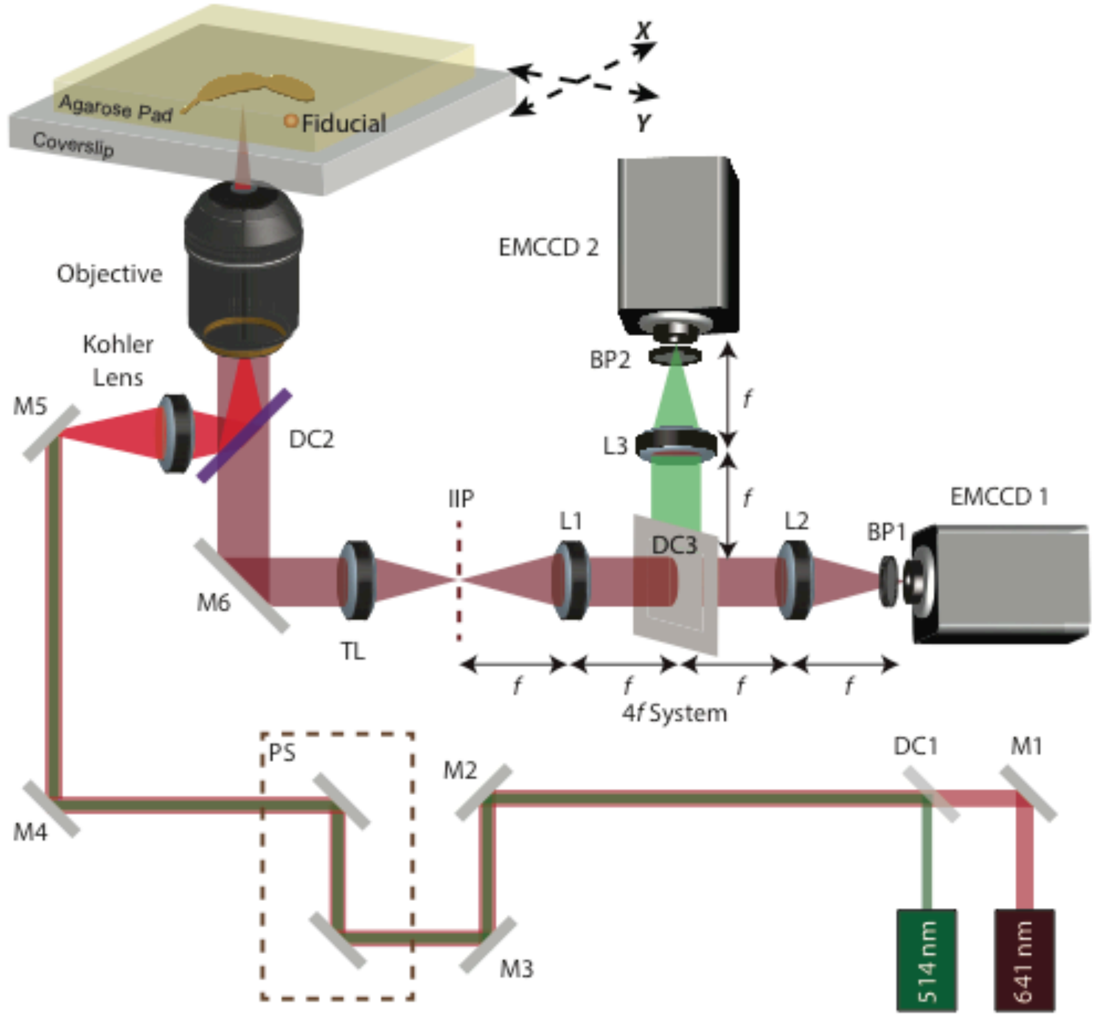


**Supp Fig 10.** 2D and 3D single molecule tracking microscope schematic. 514 and 641 nm lasers are combined using a dichroic (DC1) and directed into the microscope by a series of mirrors (M1-M5) and a periscope (PS). A Kohler lens is used to create widefield illumination. An objective focuses the lasers onto the sample, including *Caulobacter crescentus* cells (brown) and fiducial beads (orange) on an agarose pad (yellow). Fluorescence is collected through the same objective, passes through DC2, and is focused by the tube lens at the intermediate image plane (IIP). A 4f lens system (L1 & L2/L3) is used to access the Fourier plane (gray plane) where a double-helix phase mask is placed for 3D tracking experiments. The red and green detection channels are split at DC3 and are filtered by two band pass filters (BP1 & BP2) for 2-color, 2D tracking experiments. The color channels are imaged on two separate EMCCD cameras. For 3D tracking experiments, only the 641 nm laser is used and DC3 is removed.
